# Kinetics of self-assembly of inclusions due to lipid membrane thickness interactions

**DOI:** 10.1101/2020.09.23.309575

**Authors:** Xinyu Liao, Prashant K. Purohit

## Abstract

Self-assembly of proteins on lipid membranes underlies many important processes in cell biology, such as, exo- and endo-cytosis, assembly of viruses, etc. An attractive force that can cause self-assembly is mediated by membrane thickness interactions between proteins. The free energy profile associated with this attractive force is a result of the overlap of thickness deformation fields around the proteins. The thickness deformation field around proteins of various shapes can be calculated from the solution of a boundary value problem and is relatively well understood. Yet, the time scales over which self-assembly occurs has not been explored. In this paper we compute this time scale as a function of the initial distance between two inclusions by viewing their coalescence as a first passage time problem. The first passage time is computed using both Langevin dynamics and a partial differential equation, and both methods are found to be in excellent agreement. Inclusions of three different shapes are studied and it is found that for two inclusions separated by about hundred nanometers the time to coalescence is hundreds of milliseconds irrespective of shape. Our Langevin dynamics simulation of self-assembly required an efficient computation of the interaction energy of inclusions which was accomplished using a finite difference technique. The interaction energy profiles obtained using this numerical technique were in excellent agreement with those from a previously proposed semi-analytical method based on Fourier-Bessel series. The computational strategies described in this paper could potentially lead to efficient methods to explore the kinetics of self-assembly of proteins on lipid membranes.

**Author summary:** Self-assembly of proteins on lipid membranes occurs during exo- and endo-cytosis and also when viruses exit an infected cell. The forces mediating self-assembly of inclusions on membranes have therefore been of long standing interest. However, the kinetics of self-assembly has received much less attention. As a first step in discerning the kinetics, we examine the time to coalescence of two inclusions on a membrane as a function of the distance separating them. We use both Langevin dynamics simulations and a partial differential equation to compute this time scale. We predict that the time to coalescence is on the scale of hundreds of milliseconds for two inclusions separated by about hundred nanometers. The deformation moduli of the lipid membrane and the membrane tension can affect this time scale.

## Introduction

Self-assembly of proteins on lipid membranes has been a topic of interest for at least the last three decades [1–3]. Proteins on membranes self-assemble because they interact with each other through forces that have their origins in membrane bending deformations [4, 5], membrane thickness deformations [4, 6–11], electrostatics [12] and entropic interactions [4, 13]. There is a large literature on this topic that we do not attempt to review here [3–6, 13–20]. Our interest is in self-assembly caused by membrane thickness mediated interactions of proteins.

It is well known that lipid bilayers consist of two leaflets with the hydrophobic tails of the lipid molecules spanning the membrane thickness. Proteins that are embedded in the membrane have hydrophobic peptides placed in such a way as they interact mostly with the hydrophobic tails of the lipid molecules. If the thickness of the hydrophobic region of a protein is different from that of the lipid membrane then the leaflets deform so that the membrane thickness in the vicinity of the protein changes (see Fig. 1(a)). The energy cost of the thickness deformation has been estimated analytically by taking account of the lipid hydrocarbon chain entropy [9, 21]. The result is an energy functional written in terms of the deformation field *u*(*x, y*) of the half-membrane thickness and its gradients [4, 9]. The membrane bending modulus *K*_*b*_, the membrane thickness modulus *K*_*t*_ and the isotropic membrane tension *F* enter as parameters into this functional. The Euler-Lagrange equation obtained by the minimization of this energy functional is a fourth order linear partial differential equation (PDE). A series of papers by Phillips, Klug, Haselwandter and colleagues [6–8, 22] start from this energy functional and utilize the linearity of the PDE to computationally analyze allosteric interactions of clusters of proteins of various shapes. The key idea is that the thickness deformation fields caused by distant proteins can overlap (superimpose) and give rise to interaction forces just as defects in elastic solids interact due to the overlap of deformation fields [23]. This idea has been in place since at least the mid-1990s [9], but it was computationally extended to complex protein shapes and large clusters by the above authors.

**Fig 1.**
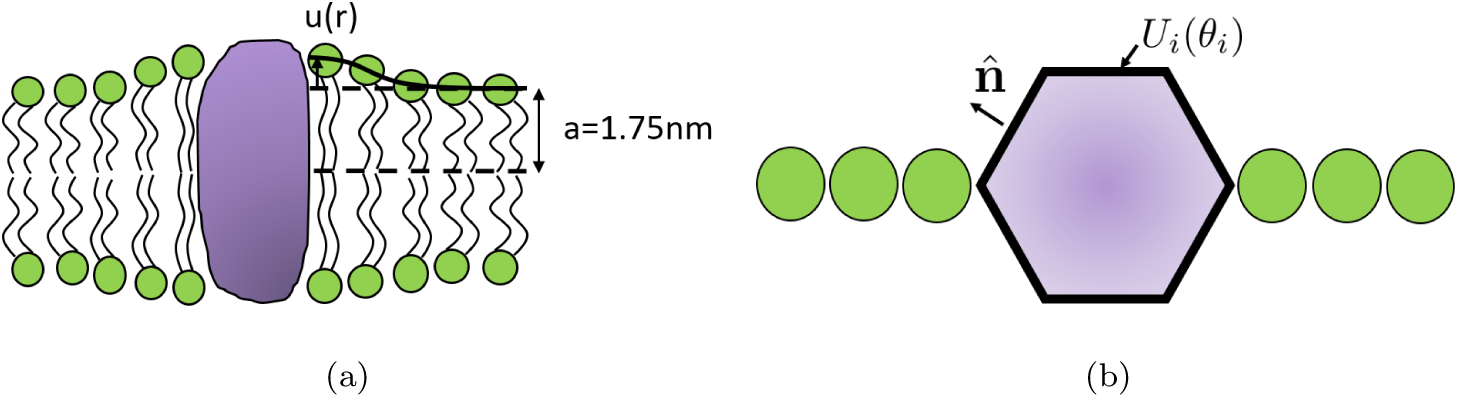
(a) Schematic of bilayer deformations due to a thickness mismatch between hydrophobic region of a bilayer leaflet and an embedded protein. (b) The two types of boundary conditions that are used in this work. Dirichlet boundary condition *U*_*i*_(*θ*_*i*_) gives the thickness deformation along the boundary of inclusion *i*, while the slope boundary condition 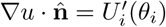 determines the derivative along normal directions at each point along the boundary of inclusion *i*. The top view of the surrounding lipid molecules (green circles) are only shown along the horizontal line, but they are everywhere on the plane.

An important result that emerged from the research on clusters discussed above is that the interaction free energy has a maximum when plotted as a function of distance between individual proteins (which form a lattice). To the left of the maximum there are strong attractive forces between the proteins, while to the right there are weak repulsive forces which decay away as the proteins move far apart. The strong attractive forces should cause self-assembly if two (or more) proteins happen to come close together as they diffuse on the membrane. We are interested in the time scale of the self-assembly process. There are few experiments which focus on this time scale, but one by Shnyrova *et al.* [24] found that viral proteins (that did not interact electrostatically) on a micron-sized vesicle self-assemble in seconds.

Temporal evolution of the self-assembly of viral proteins on a lipid membrane has been analyzed in a few recent papers using simulations. Often these simulations can be computationally prohibitive, but they do give insight about time scales and intermediate states of the cluster of proteins assembling into a virus particle or nano-container [1–3, 25]. A drawback of these simulations is that they may not be able to tackle time scales of seconds over which self-assembly was seen to occur in experiments [24]. We will take a different approach in this paper by analyzing self-assembly of differently shaped inclusions using Langevin dynamics and the corresponding Fokker-Planck equations. In recent work We viewed self-assembly of two inclusions as a first-passage time problem which can be quantitatively analyzed using the theory of stochastic processes [26]. We implemented this approach in the context of interactions based on membrane bending. The analytical calculations (using PDEs) in [26] were confined to absorbing boundary conditions on both boundaries. A novelty of this work is that we extend the PDE approach to include absorbing and reflecting boundary conditions.

This paper is arranged as follows. First, we quantify the interaction energy profile of hexagonal, rod- and star-shaped inclusions^1^. We show that our finite difference numerical method for computing energies agrees very well with analytical formulae (using Fourier-Bessel series) in most cases. After computing the interaction energies, we solve first-passage time problems to find the time scales over which two inclusions coalesce due to attractive interactions. We use both Langevin dynamics and the Fokker-Planck equation to solve first passage time problems and study both isotropic and anisotropic problems with reflecting/absorbing boundary conditions. Finally, we summarize our results in the discussion and conclusion sections and point to various enrichments that can be implemented following our earlier work [26].

**Table 1.** List of parameters

| Symbol | Description | Units | Typical values |
| --- | --- | --- | --- |
| $\ell$ | side length of triangular grid | nm | 2.5 |
| $K_b$ | bending modulus | pN·nm | 82.8 [6] |
| $T$ | temperature | K | 300 |
| $k_B$ | Boltzmann constant | N·m·K<sup>-1</sup> | $1.38 \times 10^{-23}$ |
| $K_t$ | thickness deformation modulus | pN·nm<sup>-1</sup> | 248.4 [6] |
| $r$ | separations between two inclusions | nm | 9 – 125 |
| $F$ | applied tension | pN·nm<sup>-1</sup> | 0.1 – 10 |
| $a$ | unperturbed bilayer half-thickness | nm | 1.75 [11] |
| $\mathcal{R}_1(\theta_1)$ | shape function for the centered inclusion | nm | |
| $\mathcal{R}_2(\theta_2)$ | shape function for the moving inclusion | nm | |
| $\theta$ | the angle between two inclusions and horizontal line (see Fig 2(b)) | radian/degree | |
| $u$ | thickness deformation | nm | |
| $u_i$ | thickness deformation at node $i$ | nm | |
| $A_e$ | the area of the triangular element at node $i$ | nm<sup>2</sup> | |
| $R_1$ | radius of the inner boundary w.r.t $(r_1, \theta_1)$ | nm | |
| $R_2$ | radius of the outer boundary w.r.t $(r_2, \theta_2)$ | nm | |
| $\mathbf{u}_b$ | the vector of all nodes determined by Eq (14)-(15) and inside inclusions | | |
| $\mathbf{u}_a$ | the vector of all nodes that are not in $\mathbf{u}_b$ | | |
| $\mathbf{u}$ | $\mathbf{u} = [\mathbf{u}_a^T, \mathbf{u}_b^T]^T$ | | |
| $\phi$ | Energy of the system | pN·nm | |
| $n$ | the number of refined grid elements in one side of the original triangle grid | | |
| $\nu(\nu_{ij})$ | translational drag coefficient (tensor) | s·pN·nm<sup>-1</sup> | $2.32 \times 10^{-5}$ |
| $D(D_{ij})$ | diffusion coefficient (tensor) | nm<sup>2</sup>·s<sup>-1</sup> | $1.76 \times 10^5$ |

<sup>1</sup>We are limited in the shapes we can explore by the equilateral triangle grid used in our computations.

## Energy landscape

### Analytical solution based on Fourier-Bessel basis

We consider a circular lipid membrane with radius *R*_2_ and two inclusions embedded in it. Our first goal is to compute the energy landscape seen by an inclusion interacting with another inclusion on a flat membrane. The interactions between the inclusions are a result of the overlap of membrane thickness deformation fields in their vicinity. The interaction energy will be computed by considering two inclusions, one fixed and the other moving as shown in Fig 2(a). The coordinate frame at the fixed inclusion (blue) denoted as inclusion 1 (*r*_1_, *θ*_1_) is set to be the default one. Assume that the moving inclusion (purple) denoted as inclusion 2 initially stays in the same orientation as inclusion 1 (see Fig 2(a)). To keep the analysis simple, when an inclusion moves we do not consider its rotational diffusion. As inclusion 2 moves from the initial configuration to the green spot and forms an angle *θ* with the horizontal line (see Fig 2(b)), the energy of the system can be computed by rotating both inclusions anticlockwise by angle *θ* from the initial configuration (see Fig 2(c)). This interaction energy will enter our analysis of the kinetics of the moving inclusion due to Brownian motion.

**Fig 2.**
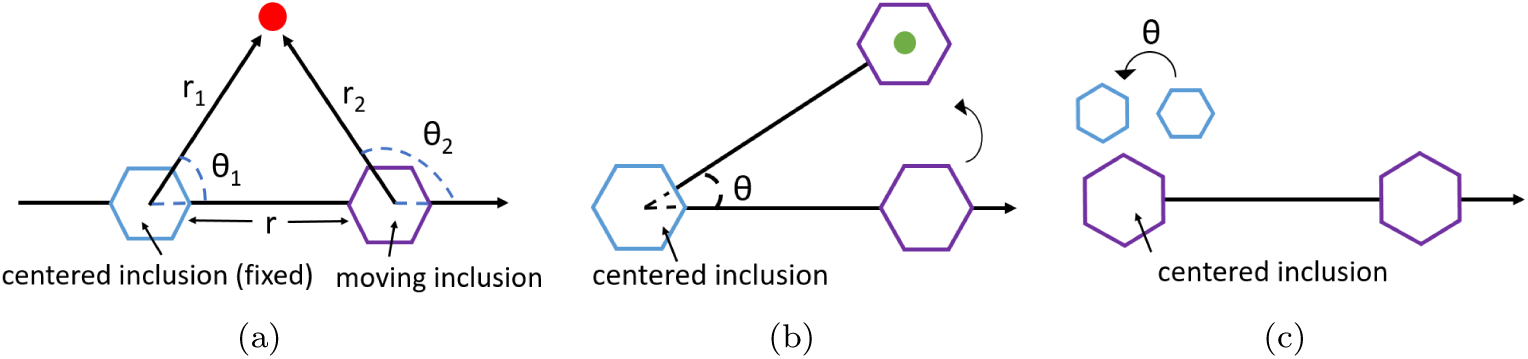
(a) The initial configuration of a system of two inclusions. The fixed inclusion located at the center (blue) has local coordinate: (*r*_1_, *θ*_1_) and the moving inclusion (purple) has local coordinate: (*r*_2_, *θ*_2_). (b) The inclusion on the right moves to the green spot and forms an angle *θ* with the horizontal line. (c) The energy of the configuration here is the same as the one in (b). Note that the hexagons in (c) are rotated when compared to hexagons in (a).

The elastic energy due to thickness deformation is given by [7, 8, 10, 27],

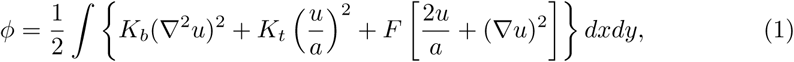

The Euler-Lagrange equation associated with Eq (1) is given by [7],

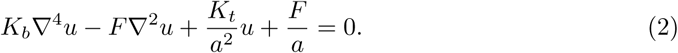

Eq (2) can be reduced to the following form using the transformation 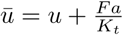,

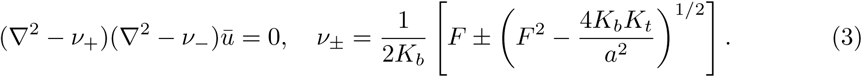

First, we consider the case of an infinitely large circular membrane with *R*_2_*→ ∞* without applied tension (*F* = 0). We assume natural boundary condition which means that *u* = *ū →* 0 at *∞*. Let inclusion 2 be on the right side of inclusion 1. Then, a Fourier-Bessel series solution for the thickness deformation field around each inclusion *i*(*i* = 1, 2) can be obtained,

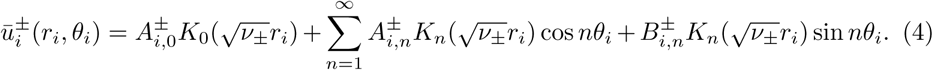

For small applied tension *F* and large membrane size *R*_2_, we used Eq (4) as an approximation for the solution of 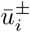. Since the Euler-Lagrange equation (Eq (2)) is linear, the solution for Eq (3) is given by [8],

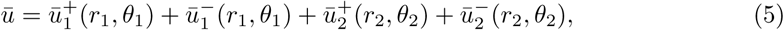

in which we used the coordinate transformations,

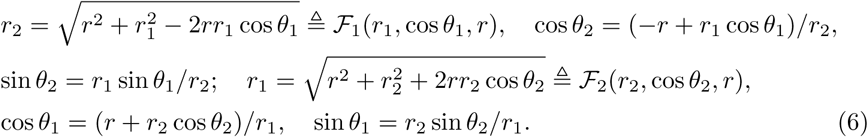

In order to efficiently apply the boundary conditions, we rewrite *ū*_2_ as a function of *r*_1_, *θ*_1_, *r* and *ū*_1_ as a function of *r*_2_, *θ*_2_, *r*,

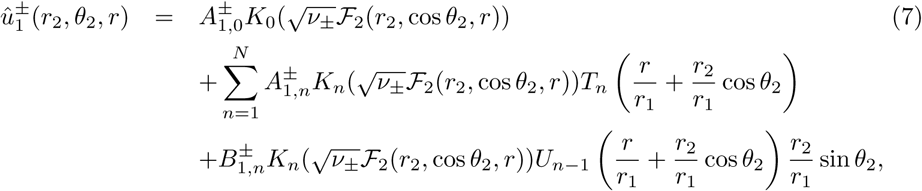

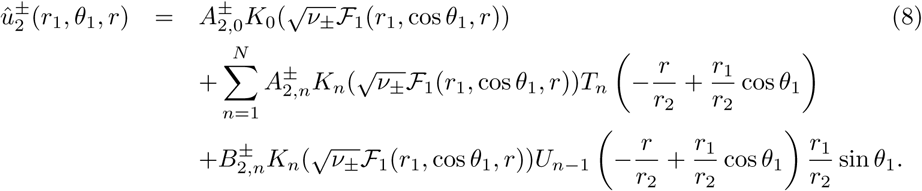

Let 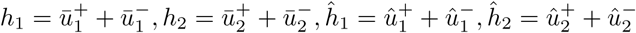. We consider the following type of boundary conditions (see Fig 1(b)),

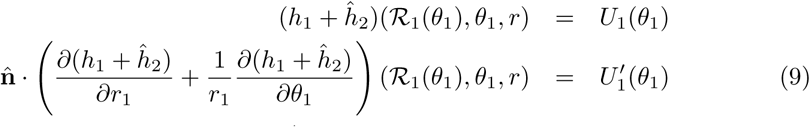

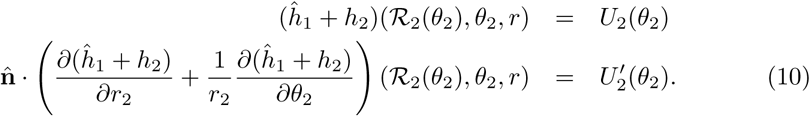

We can solve for the 4(2*N* + 1) coefficients 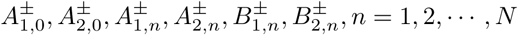 because Eq (9)-(10) result in a linear system. This determines the full deformation field due to the overlap of the deformations caused by both inclusions. The next step is to compute the energy *ϕ*(**r**) due to this deformation field. Note that the angular dependence of *ϕ*(**r**) appears in the shape functions of two inclusions, ℛ_1_, *ℛ*_2_.

Using the divergence theorem, the total energy expression in Eq (1) can be converted to the sum of line integrals over the boundary terms, i.e. *φ* = *ϕ*_1_ + *ϕ*_2_ with *ϕ*_*i*_ given by,

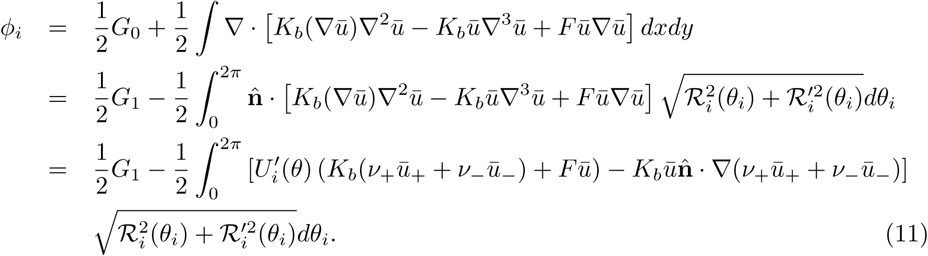

From the first line to the second line we assume the line integral along the outer boundary is a constant (which works out to 0 as *R*_2_ *→ ∞* and *F →* 0) w.r.t *r* and put it into the *G*_1_ term (both *G*_0_ and *G*_1_ are constants). The energy *φ*(**r**) can be computed relatively efficiently using this technique. This is important since *φ*(**r**) must be computed repeatedly as inclusion 2 moves and **r** changes due to Brownian motion when we solve the first passage time problem. We will also need the forces acting on inclusion 2 in our analysis later. Eq (11) gives an expression to compute the force analytically, which in the special case of an isotropic *φ*(**r**) (i.e., no angular dependence) works out to

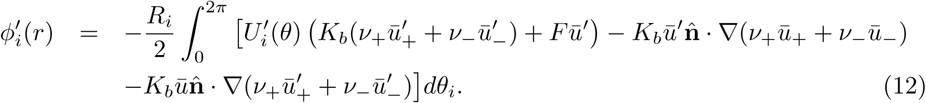

When there is only one circular inclusion in the membrane, the thickness deformation field in Eq (5) has a closed form solution [7] which can be compared to the simulation result of Klingelhoefer et al. [28] who studied radial bilayer thickness profiles for the *Gα* nanopore (among many others). We used the same parameters and boundary conditions as they did: 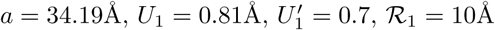 and fit their curves by choosing *K*_*t*_ = 120pN ·nm^*−*1^ and *K*_*b*_ = 2pN ·nm. The agreement shown in Fig 3 justifies the analytical method used here.

**Fig 3.**
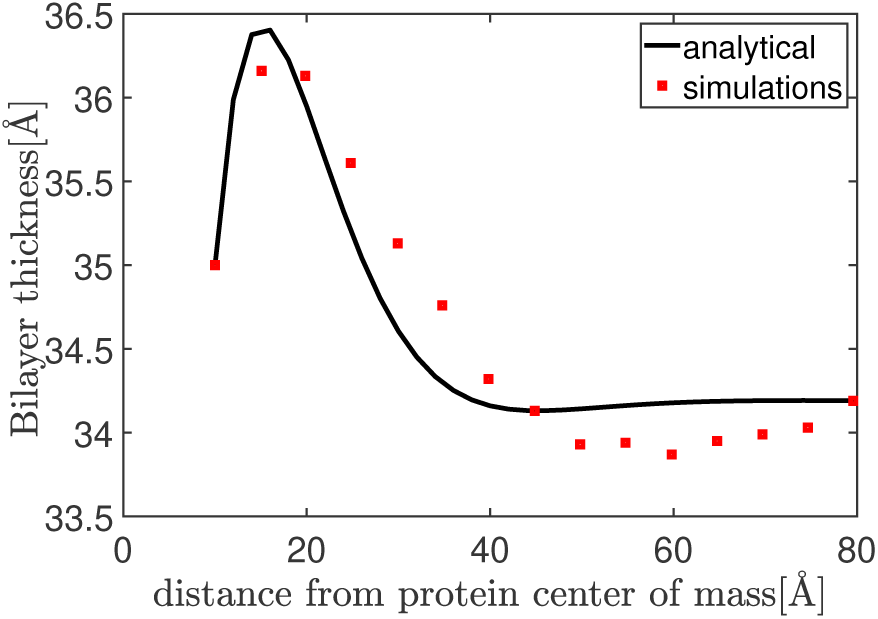
Red squares are data from the simulation by Klingelhoefer et al. [28] and the black curve is fitted using an analytical solution. The good agreement of the two profiles suggests that the energy functional Eq (1) and the associated Euler-Lagrange equation are a good starting point for estimating interaction energies of inclusions.

### Finite difference method based on refined grid

The above analysis gives us a semi-analytical technique to compute *φ*(**r**). However, not all problems can be solved analytically, so a computational method is needed to estimate interaction energies. Fortunately, Eq (1) can be minimized using a finite difference method. We discretize the membrane using equilateral triangle elements as in [29, 30]. The thickness deformation at node *i* is denoted by *u*_*i*_. Since the thickness deformation *u* changes rapidly in the neighborhood of the two inclusions, we used refined grids in a region containing the two inclusions and non-refined grids in the region far away (see Fig 4(b)) to get accurate solutions with low computational costs. Based on the triangular grids used, we study three types of inclusions shown in Fig 4(a).

**Fig 4.**
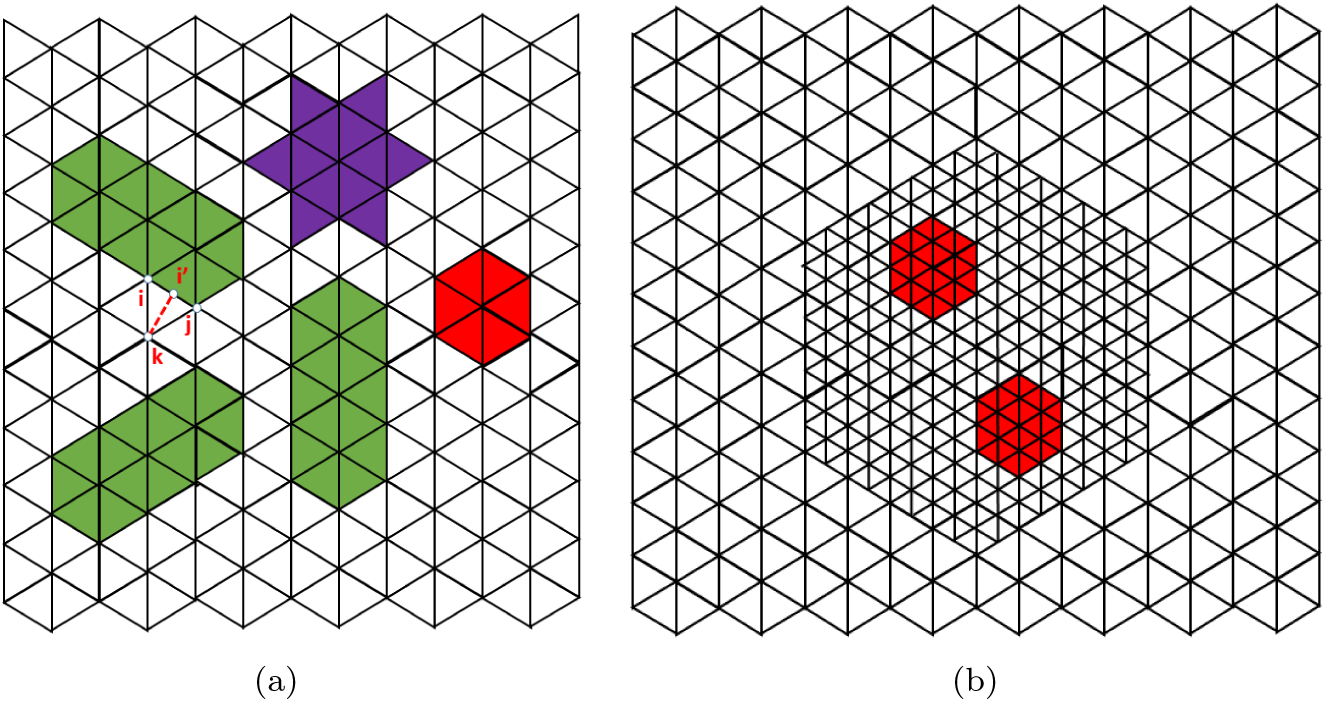
(a) Three types of inclusions studied in this paper: hexagon(red), star(purple), rod(green). (b) Refined grids are implemented in a region containing two inclusions. Here *n* = 2.

Using methods similar to those in [29, 30] the energy is first written in a discrete form and then the thickness deformation field that minimizes this energy is computed. Finally, the minimizer is plugged back into the energy expression, and we have,

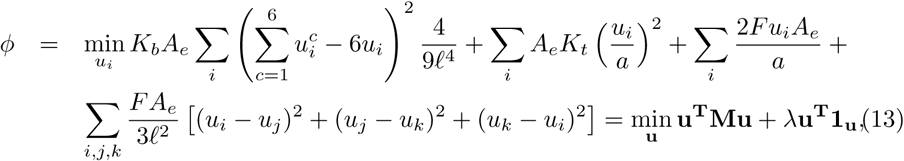

where **1**_**u**_ is a column vector of size len(**u**) with all entries 1 and 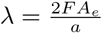. It can be shown that the boundary condition in [22] can be written in the discrete form,

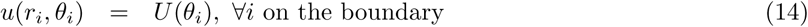

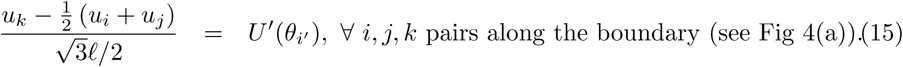

Note that *u*_*i*_, *u*_*j*_ are given in Eq (14) and thus *u*_*k*_ can be sloved from Eq (15) immediately. We also assume the inclusions are stiff such that for all nodes *i* inside the inclusions *u*_*i*_ are a constant (equal to the value of those at boundary nodes). Hence, Eq (13) can be rewritten as,

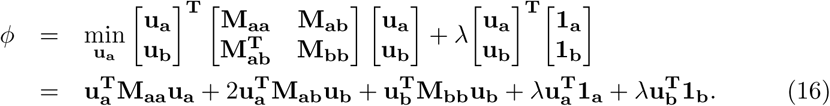

Taking 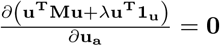, we get 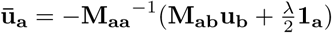 at which Eq (16) is minimized where **1**_**a**_ is a column vector of size len(**u**_**a**_) with all entries 1. Then, we can write the minimized total energy as,

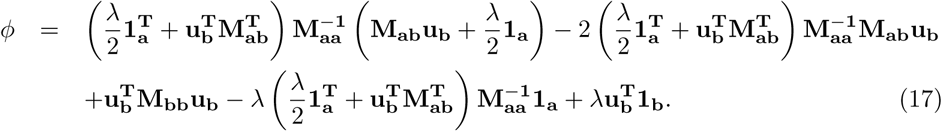

### Applications to hexagon, rod and star shaped inclusions

We now focus on the interaction of two hexagon shaped inclusions on a lipid membrane which has a rotational periodicity of *π/*3. We use Eq (17) derived from our numerical method and Eq (11) derived from the analytical method, to compute the interaction energy of two inclusions with distance *r* and then make comparisons. Agreement of the results from the two methods is expected. As shown in Fig 5(a), the energy computed using the analytical method for two inclusions separated by distance *r* in two different orientations (shown in the inset differing by a rotation of *π/*6) are almost the same. Hence, we can simplify our model and consider the energy landscape generated by two hexagon inclusions as being almost isotropic (insensitive to rotation). In Fig 5(b), we fix the shape of two hexagons (see inset of the figure), and show that as the number of grid elements per side *n* increases, the match between the energy computed from the numerical method and analytical method gets better, justifying our numerical approach of using refined grids near the inclusions. From Fig 5(c) we learn that as applied tension increases, the attraction at small separations (around *r* = *R*_1_ = 7 nm) becomes weaker but the repulsive force at around 9 *−* 10nm also becomes weaker. Please check those to be sure.

**Fig 5.**
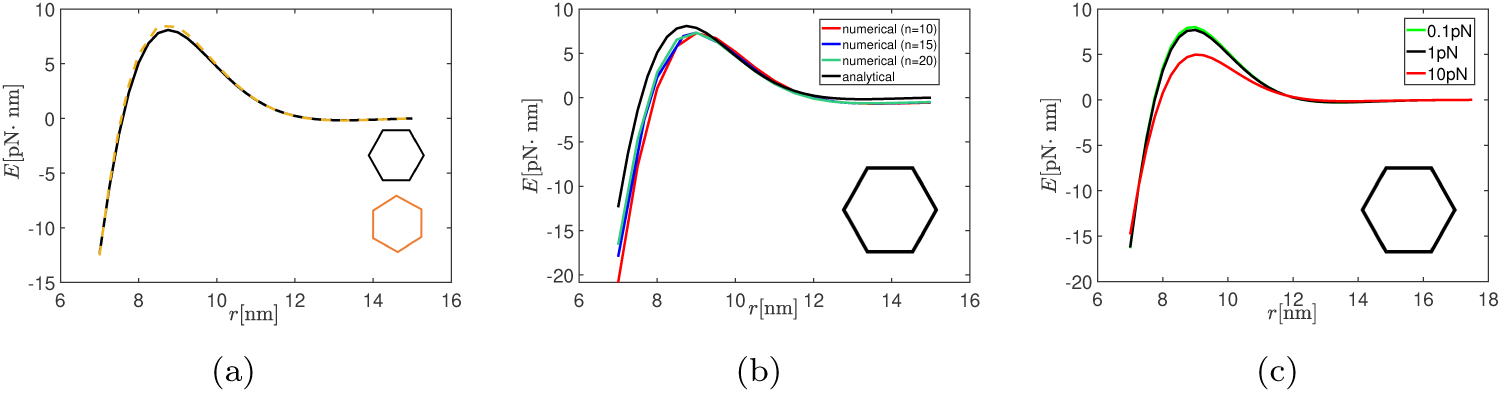
(a) The energy computed by analytical method using Eq (11) between two configurations of hexagon inclusions up to a rotation by *π/*6 under *F* = 1pN ·nm^*−*1^. (b) The energy of the configuration under *F* = 1pN ·nm^*−*1^ computed numerically using Eq (17) converged to the energy computed by Eq (11) as the number of elements increases. (c) A comparison of the energy profiles for three different applied tension: 0.1pN·nm^*−*1^, 1pN·nm^*−*1^, 10pN·nm^*−*1^.

Fig 6(a) shows that the energy landscape of two rod shaped inclusions is anisotropic - at small separations the force is repulsive at *θ* = 0° and becomes attractive at some angle around 40° *< θ <* 50°. The attraction increases as *θ* goes up to 90°. This behavior of the energy of two rod shaped inclusions is reminiscent of the energy from out-of-plane deflection for two rods [26] on a lipid membrane. Fig 6(b) shows that the energy computed by the numerical method and analytical method again agree very well which gives us confidence in the numerical method.

**Fig 6.**
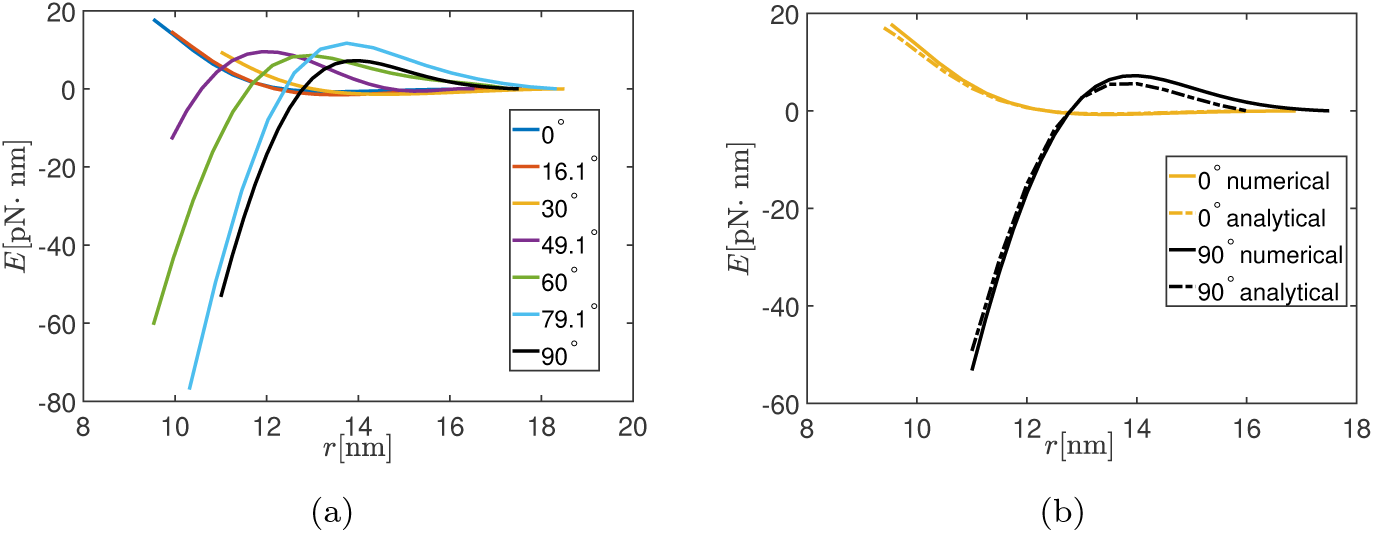
(a) The energy of two shaped rod inclusions with different *θ* under applied tension 1pN ·nm^*−*1^. (b) The energy computed by analytical method using Eq (11) fits the one computed by numerical method using Eq (17) both with *θ* = 0° and *θ* = 90° under applied tension 1pN·nm^*−*1^

Next we compute the interaction energy of rod shaped inclusions for tensions *F* = 0.1pN ·nm^*−*1^ and *F* = 10pN· nm^*−*1^. The comparison between Fig 7 and Fig 6(a) shows that as applied tension increases, the force becomes weaker at short separations, which implies that elastic interactions could be weakened by strong applied tension. Physically, this is reasonable, since high tension will tend to make the membrane flatter so that the thickness is more uniform everywhere.

**Fig 7.**
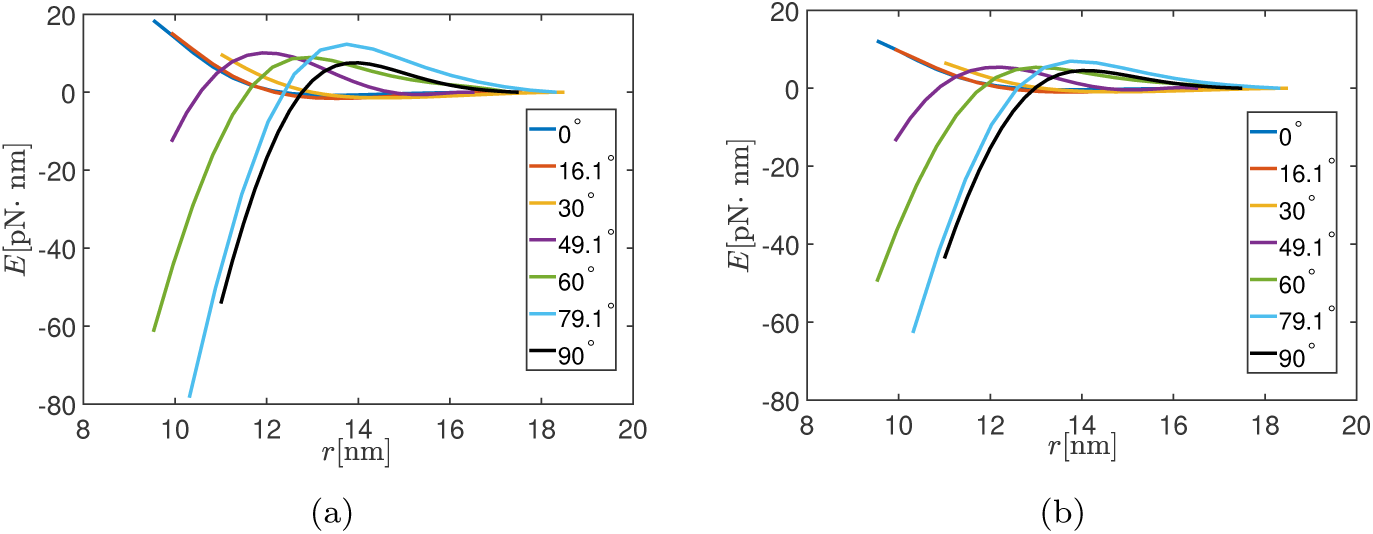
(a) The energy of two rod shape inclusions under different *θ* with applied tension 0.1pN·nm^*−*1^. (b) The energy of two rod shape inclusions under different *θ* with applied tension 10pN·nm^*−*1^.

Next, we apply both methods to compute the interaction energy of two star shaped inclusions in Fig 8. Just as in the case of hexagons, we consider various orientations of the star shaped inclusions as shown in the inset of Fig 8(a). The match between the analytical method and numerical method is not as good in this case because the star shaped inclusion has 12 vertices at which the derivative along normal directions are discontinuous. Since in the analytical method we used Fourier-Bessel series to approximate the contour (ℛ_1_, *ℛ*_2_) and the derivative along normal directions to the boundaries, it requires a large number of terms *N* to obtain a good approximation. This is computationally not feasible for symbolic operations in MATLAB. Thus, we have greater confidence in our finite difference numerical method to compute interaction energies in complex geometries. In Fig 8(b) we use our numerical method to compute the interaction energies for star shaped inclusions for various values of *F*. The trends are similar to those seen for hexagon shaped inclusions.

**Fig 8.**
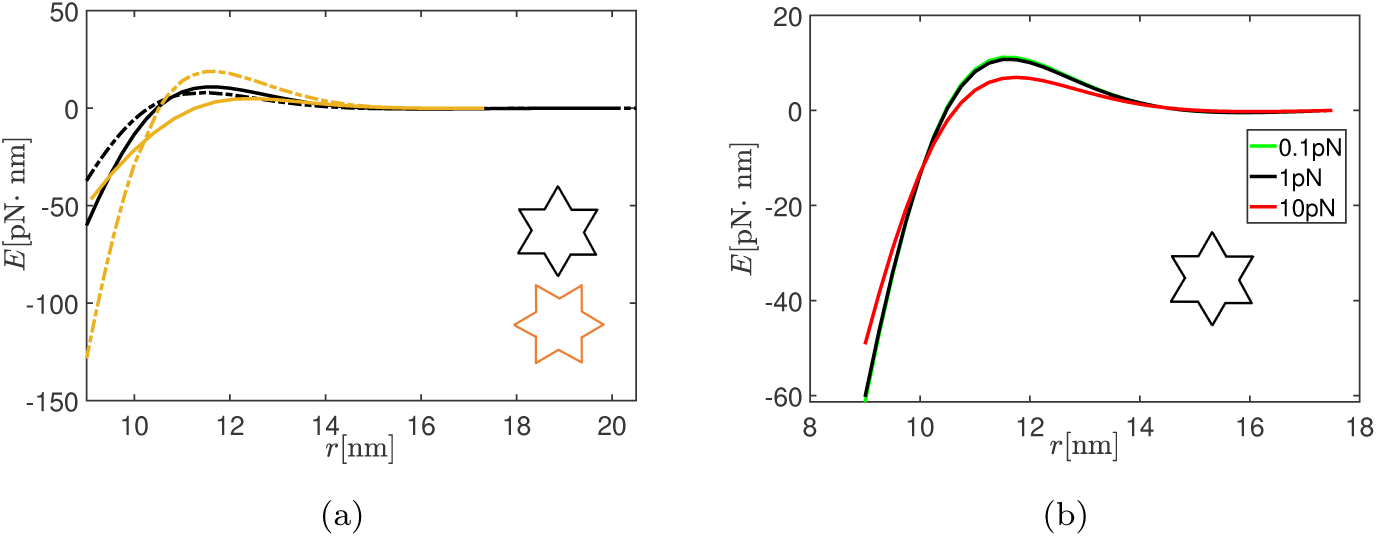
(a) Solid lines are the energy computed by numerical method using Eq (17) and dashed lines are the energy computed by analytical method using Eq (11). The applied tension is 1pN·nm^*−*1^. (b) The energy computed by numerical method under three applied tensions: 0.1pN·nm^*−*1^, 1pN·nm^*−*1^, 10pN·nm^*−*1^.

## First passage time for isotropic inclusions under mixed boundary condition

Our main goal in this paper is to study the kinetics of an inclusion diffusing in an energy landscape resulting from elastic interactions with another inclusion. Efficient methods to compute the energy landscape developed above are a pre-requisite to this exercise. We will now use these methods to solve first passage time problems.

We consider a circular membrane of size *R*_2_ = 125 nm with a circular inclusion of size 2.5 nm fixed at the center. Another circular inclusion of the same size is diffusing around driven by stochastic forces. Recall from the energy landscape that there are attractive interactions between inclusions when the separations are small. Hence, if the moving inclusion comes close enough to the static one at the center then it will be strongly attracted. Therefore, we assume that at *R*_1_ = 7 nm there is an absorbing wall at which the moving inclusion will disappear by being attracted towards the center. We assume that at *R*_2_ = 125 nm (far away) there is a reflecting wall where the moving inclusion will be bounced back. Note that problems in which both boundaries are absorbing were solved elsewhere [26]. The exercise we will perform now is as follows. We place the second inclusion randomly on a circle of radius *r* = *y* at time *t* = 0 and let it diffuse around. At some time *t* = *T*_*in*_ when the inclusion hits the inner boundary for the first time we stop it from diffusing and record *T*_*in*_. We repeat this experiment a large number of times and record *T*_*in*_ for each repetition. The mean value of *T*_*in*_ is the mean first passage time *T*_1_. Our goal is to find *T*_1_(*y*) as a function of the initial condition *r* = *y*. This can be done analytically or through a Langevin dynamics simulation. We will use both methods in the following.

To estimate *T*_1_(*y*) analytically we first need to compute survival probabilities. Let *p* be the probability density (for finding the inclusion) at position *r* and angle *θ* given initial condition *r* = *y, θ* = *α* and 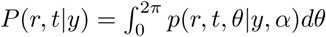. The probability density *p* is independent of *θ* since neither the energy landscape nor the diffusion (or drag) coefficient of the inclusion depends on it. As a result, the Fokker-Planck equation for the evolution of this probability is in the following isotropic form [26],

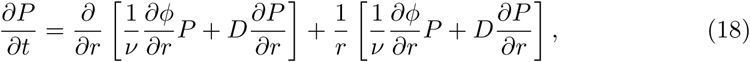

with Dirichlet boundary condition at the inner boundary and Robin boundary condition at the outer boundary [31],

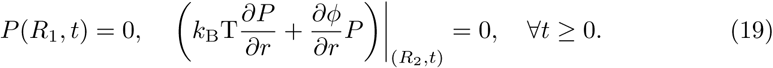

In the above *D* is a diffusion coefficient of the inclusion in the lipid membrane and *ν* is a drag coefficient which are related by the Nernst-Einstein relation *νD* = *k*_B_T [26]. Let *S*(*y, t*) be the survival probability,

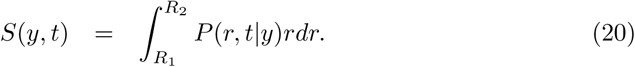

Then, we can get the first passage time density,

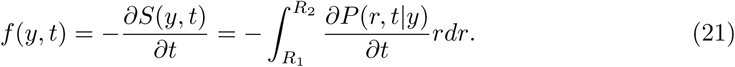

The existence of the first moment of *P* (*r, t|y*) with respect to time *t* can be shown from the fundamental solution constructed by Itô in [32]. Then, *tP* (*r, t|y*) *→* 0 as *t → ∞*. Accordingly, the first passage time *T*_1_(*y*) can be derived from Eq (21),

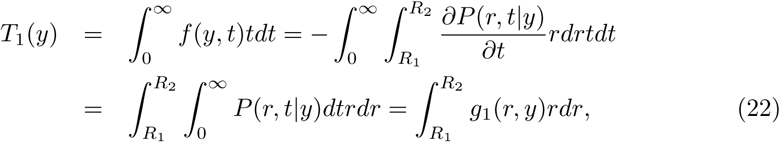

where *g*_1_ is defined by,

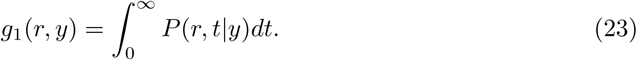

### Theorem 1

The ODE for *T*_1_(*y*) with a reflecting wall at the outer boundary and an absorbing wall at the inner boundary is

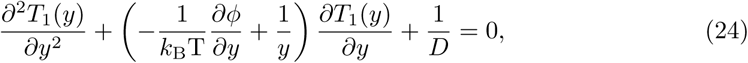

with boundary conditions,

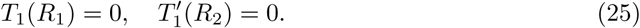

*Proof:* See Proof of Theorem 1.

Next, we describe how to estimate *T*_1_(*y*) using Langevin dynamics simulations. The overdamped version of the Langevin equation in an isotropic setting is given by [26],

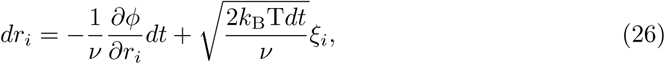

where *i* represents two perpendicular directions of the motion. *ν* is the translational drag coefficient of a circular inclusion given by the Saffman-Delbrck model [33]. *ξ*_*i*_*∼ 𝒩* (0, 1), a normally distributed random variable with mean 0 and variance 1, represents the stochastic force along direction *i*. We initially put the moving particle somewhere at *r* = *y*, and choose a time step *dt* that ensures convergence of the Lagenvin dynamics simulation. Then, for each time step *dt*, we perform the calculation in Eq (26), updating the position of the moving inclusion. We record the time at which the moving particle hits the absorbing wall at *R*_1_. We run 8000 simulations and then take an average to estimate the first passage time. For more details the readers are referred to [26]. Fig 5(a) and Fig 8(a) show that the *φ*(**r**) for hexagon and star inclusions can be regarded as nearly isotropic. We use Eq (24) to numerically solve for the first passage time and compare the results obtained from the two methods.

Fig 9 shows that the first passage time for hexagonal inclusion derived from the two methods are in good agreement. As the applied tension increases, the first passage time is reduced at most *r* that are not close to *R*_1_. At first glance this might seem counter-intuitive because from Fig 5(c) we know that at small separations (close to *R*_1_) the attraction force becomes weaker as applied tension increases. However, there is a stronger repulsive force at around *r* = 9 *−* 10nm under large applied tension which slows the motion of the moving particle from a large starting separation.

**Fig 9.**
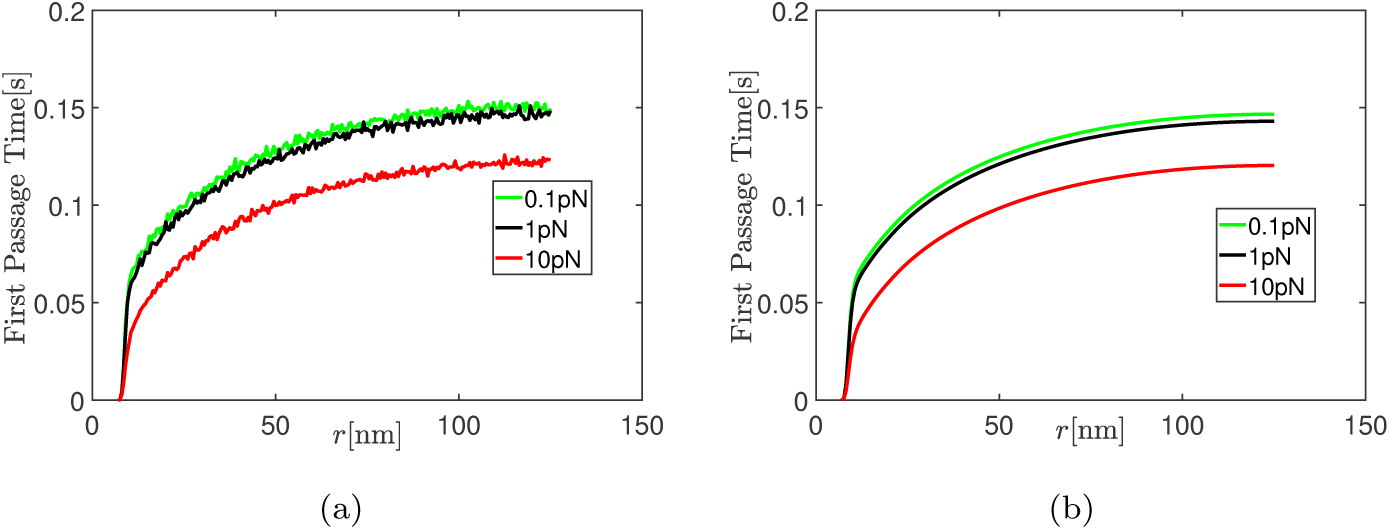
The first passage time for hexagon shaped inclusions is computed using (a) Langevin dynamics simulations in Eq (26), (b) ODE in Eq (24) under three applied tensions 0.1pN·nm^*−*1^, 1pN·nm^*−*1^, 10pN·nm^*−*1^.

The first passage time computed by the two methods is also in good agreement when the inclusions are star shaped. The order of the first passage time is the same as the hexagonal inclusions and similar arguments for the shape of the curves can be made here.

## First passage time for anisotropic inclusions under mixed boundary condition

For two non-circular inclusions, the corresponding Fokker-Planck equation is a partial differential equation of parabolic type [26],

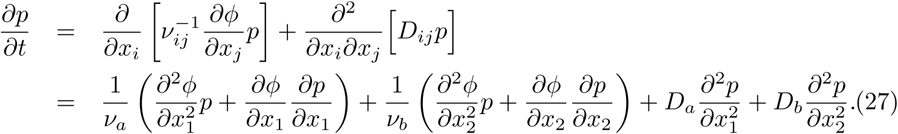

Accordingly, we need to redefine the first passage time in Eq (22) which is now given by

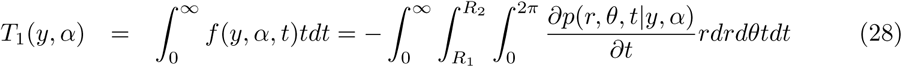

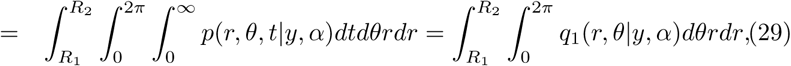

where *tp*(*r, θ, t|y, α*) *→* 0 as *t → ∞* is implemented in the first equation of the second line and *q*_1_ is defined by,

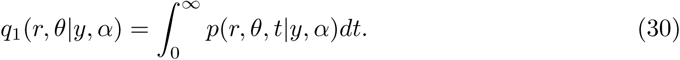

### Theorem 2

The PDE for *T*_1_(*y, α*) with a reflecting wall at the outer boundary and an absorbing wall at the inner boundary is given below,

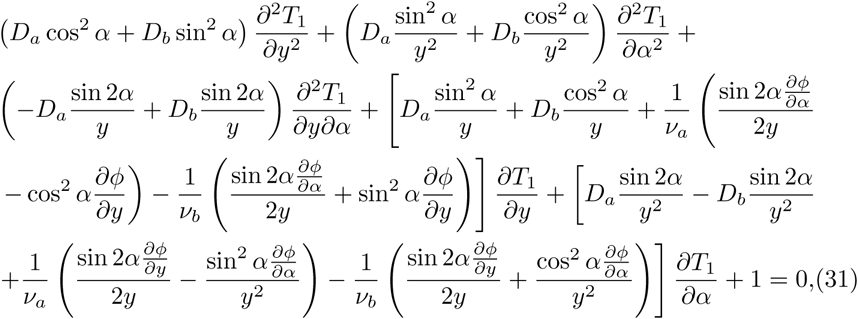

with boundary conditions

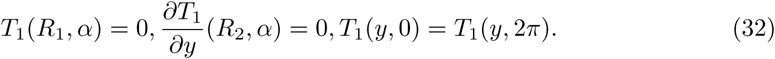

*Proof:* See Proof of Theorem 2.

The overdamped Langevin equation in an anisotropic setting is given by [26],

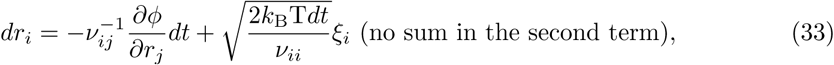

where *i* represents two perpendicular directions of the motion and *ν*_*ij*_ and *ξ*_*i*_ are the diffusion tensor and random force tensor (for more details see [26]). Fig 6(a) shows that the interaction energy *φ*(**r**) for rod shaped inclusions depends on *θ* (it is anisotropic). In the Langevin dynamics calculations, for each initial position *y* we use Eq (33) to run 8000 simulations with a reflecting wall at *R*_2_ and an absorbing wall at *R*_1_ for four *θ* = 0°, 30°, 60°, 90° and then take an average (for each *θ* separately) to estimate the first passage time. We also use Eq (31) to numerically solve the first passage time and compare the results derived from the two methods for *F* = 0.1, 1, 10 pN/nm.

The good agreement between the first passage time solved from the PDE in Eq (31) and estimated by Langevin equation once again shows that our methods are work well. As shown in Fig 11, as the initial angle increases from 0° to 90°, the first passage time decreases at small separations but increases at large separations. This can be similarly explained by the fact that stronger attractive force near *R*_1_ pulls the moving particle to be absorbed faster from smaller initial separations while stronger repulsive force around 12 *−* 16 nm leads to a larger first passage time when the particle is initially located at a large distance.

**Fig 10.**
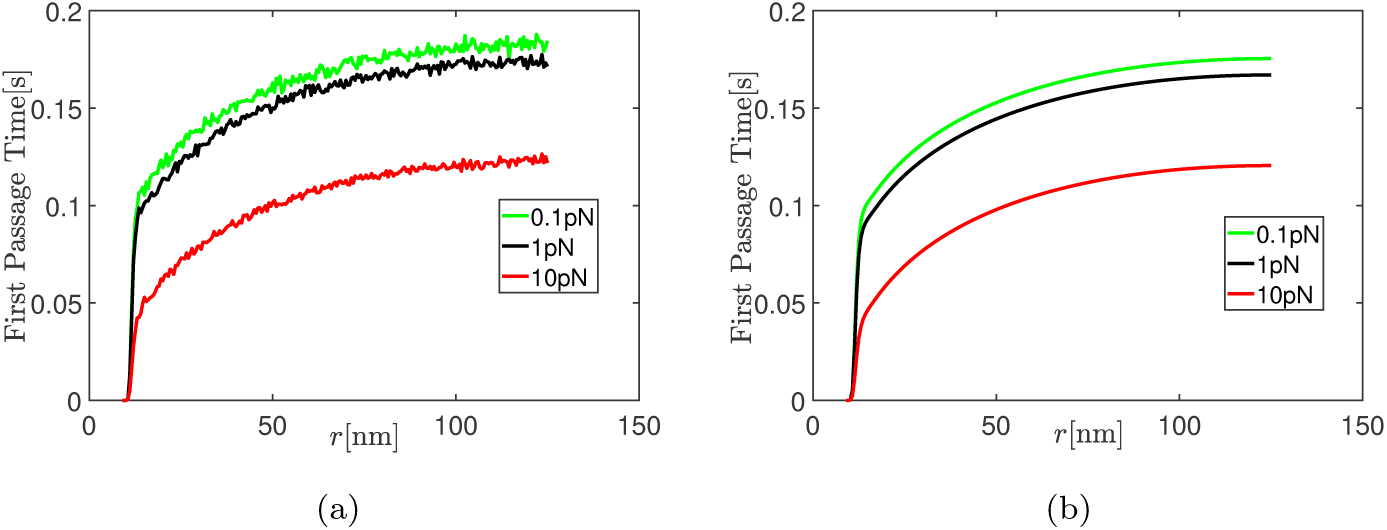
The first passage time for star-shaped inclusions are computed by (a) Langevin dynamics simulations in Eq (26), (b) ODE in Eq (24) under three applied tensions 0.1pN·nm^*−*1^, 1pN·nm^*−*1^, 10pN·nm^*−*1^.

**Fig 11.**
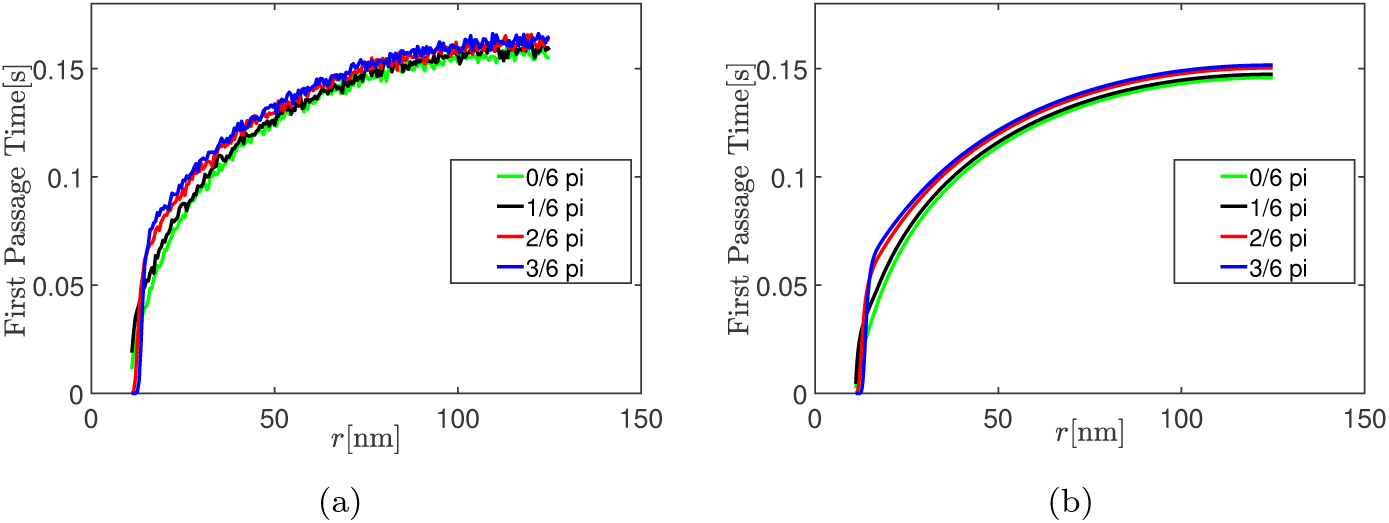
The first passage time for rod-shaped inclusions computed from (a) Langevin dynamics using Eq (33), (b) PDE using Eq (31).

The result of the first passage time under 1pN/nm applied tension in Fig 12 is similar to the one under 0.1pN/nm applied tension.

**Fig 12.**
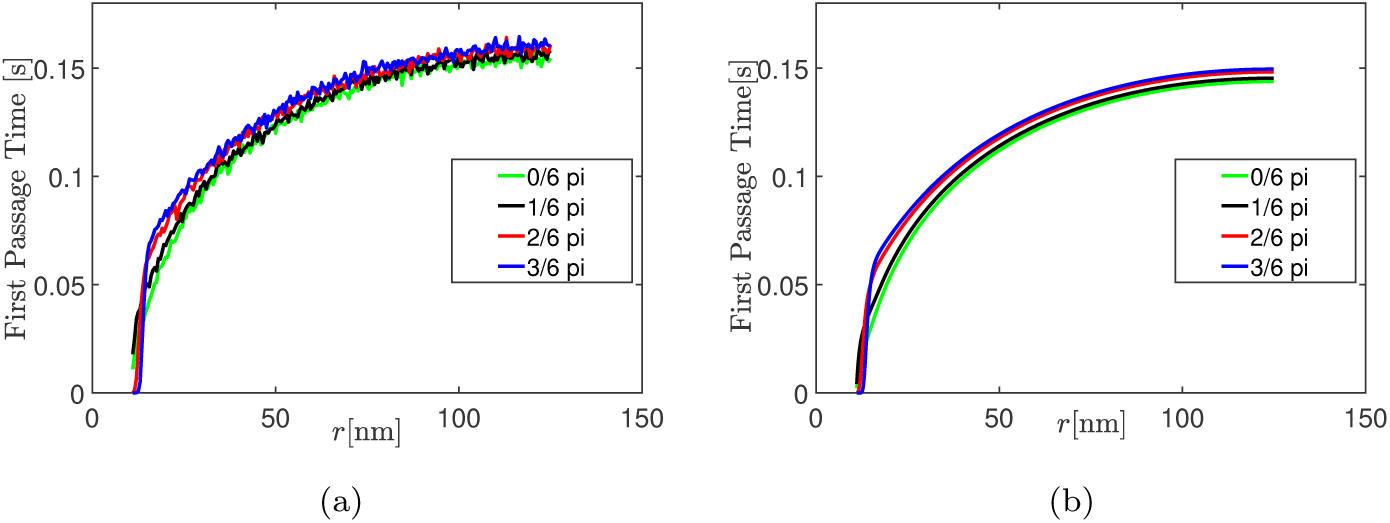
The first passage time for rod-shaped inclusions computed from (a) Langevin dynamics using Eq (33), (b) PDE using Eq (31).

**Fig 13.**
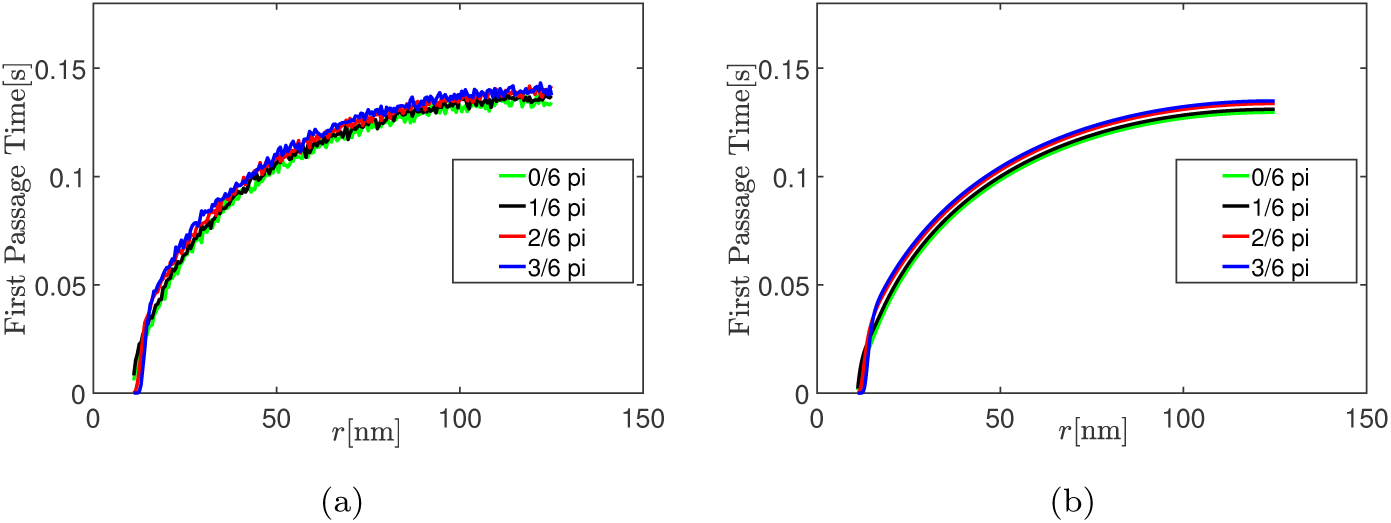
The first passage time computed from (a) Langevin dynamics using Eq (33), (b) PDE using Eq (31).

Compared to the results under 0.1,1pN ·nm^*−*1^ applied tension, the first passage time is reduced under 10pN ·nm^*−*1^ applied tension. The order of magnitude of the first passage time under all three tensions is similar.

## Discussion

This paper has two major parts. In the first part we use a finite difference method to compute the interaction energy of two inclusions due to membrane thickness deformations. In the second part we use the computed energy landscape to solve first passage time problems. Our method to compute energies is different from the analytical method in [7, 8] which uses perturbation theory to study thickness mediated interactions between two anisotropic inclusions; we implement an approach to compute the energy using the divergence theorem which is more general and can deal with strongly anisotropic inclusions. The advantage of analytical methods in both [7, 8] and this work is that they can compute the energy accurately at small applied tension *F* if enough terms in the Fourier-Bessel series are used. However, it is time consuming to compute the coefficients in the Fourier-Bessel series and this becomes computationally infeasible when the inclusions are strongly anisotropic. On the other hand, our numerical method is able to handle arbitrary values of *F* and can efficiently compute the interaction energy of two inclusions for different separations **r** given a fixed set of parameters (*K*_*b*_, *K*_*t*_, *a* etc.) which are stored in a pre-calculated stiffness matrix.

In the second part of the paper we compute the time to coalescence of two inclusions of various shapes as a function of the distance separating them. We use both Langevin dynamics and a PDE to arrive at our estimates. For two inclusions separated by about 125 nm we predict that the time to coalescence is hundreds of milliseconds irrespective of the shape of the inclusion. The time to coalescence with only membrane bending interactions was of similar magnitude as shown in [26]. The order of magnitude of the time to coalescence is the same even though the attractive force due to membrane thickness interactions is stronger than that due to membrane bending interactions in [26] at small separations. The reason is that even with membrane thickness interactions the attractive force decays to zero quickly and Brownian motion dominates the kinetics of the moving particle in most regions, just as in [26]. Therefore, at small separations the first passage time with thickness mediated interactions is smaller than that with out-of-plane bending interactions, but is not very different at large separations.

## Conclusion

In this paper we have analyzed the temporal self-assembly of inclusions due to interactions mediated by membrane thickness variations. A major accomplishment of the paper is to show that the results from Langevin dynamics simulations agree well with those obtained from a PDE for the first passage time. The approach based on the PDE is much faster than the Langevin dynamics simulation and could open new ways to study the process of self-assembly. This is a step beyond earlier studies which focused on the energy landscape of clusters of proteins, but did not look into kinetics. Some papers based on molecular simulation did consider the temporal process, but to the best of our knowledge most did not reach the time scales calculated in this paper. We close this paper by mentioning some effects that we did not consider. First, hydrodynamic interactions between inclusions (based on the Oseen tensor) were shown to speed up self-assembly in [26] and they are expected to have a similar effect here. Second, the temporal behavior of a cluster of inclusions are not studied in this paper due to limitations of computational power, but we expect the overall behavior to be similar to the clusters studied in our earlier work [26]. Third, only a limited set of inclusion shapes are considered in this paper, but it is found that the time to coalescence does not depend strongly on shape. We leave it to future work to add these refinements and extend this type of analysis to important functional proteins such as ion-channels [22].

## Supporting information

**Proof of Theorem 1** Following techniques in [26, 31, 34], we integrate Eq (18) for *P* over all *t ≥* 0,

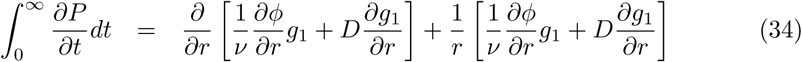

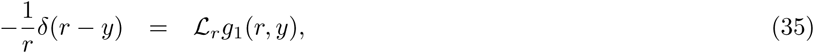

where 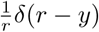 is the initial condition and the second order linear differential operator *ℒ*_*r*_ : *𝒟*(*ℒ*_*r*_) *⊂ C*^2^([*R*_1_, *R*_2_]) *→ C*^2^([*R*_1_, *R*_2_]) is defined as,

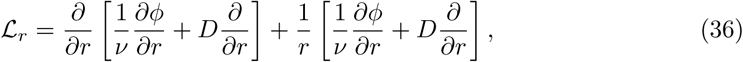

with domain

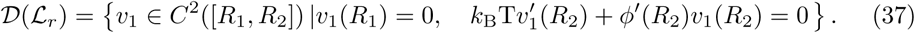

Using the method in [26], we can get the adjoint operator 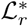 which satisfies 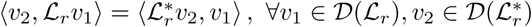,

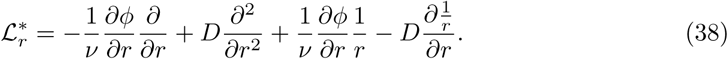

with domain

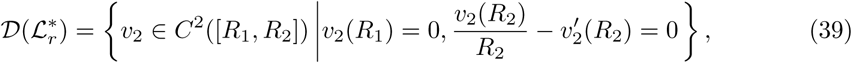

and the inner product is defined as,

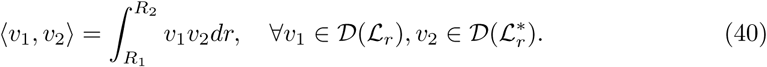

Proofs for the existence of the solutions of second order inhomogeneous linear ordinary differential equation are well known. Hence, we can find a *u*_0_ *∈ C*^2^([*R*_1_, *R*_2_]) s.t. 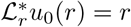. Then, it follows from Eq (22) that,

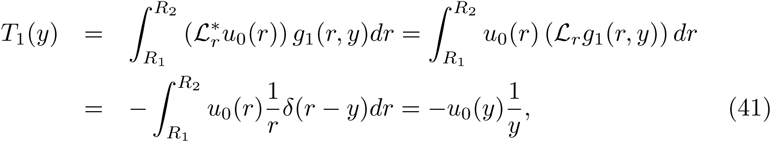

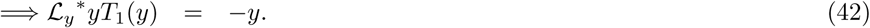

Using Eq (38), we can derive Eq (24), a second order ODE for *T*_1_(*y*). The boundary condition of *T*_1_(*y*) at the absorbing wall is straightforward [31, 34]: *T*_1_(*R*_1_) = 0. For the boundary condition at the reflecting wall, we appeal to the Langevin equation in Eq (26). If the particle sits at position *R*_2_, decomposing the overdamped Langevin equation [26] into radial direction and angular direction, we have,

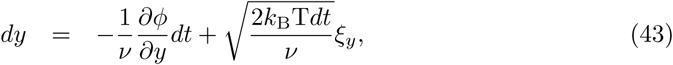

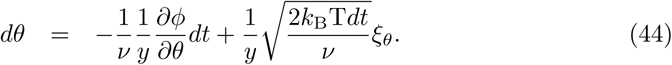

After time *dt*, the particle can only move to *R*_2_ + *dy*(*dy <* 0) along the radial direction because of the reflecting wall at *R*_2_. The motion along the angular direction can be neglected because *T*_1_(*y*) does not have dependence on angular direction. Note that *dy* is a random variable depending on *ξ*_*y*_ and *dt* with constraint *R*_1_ *≤ R*_2_ + *dy ≤ R*_2_. Then, we can write

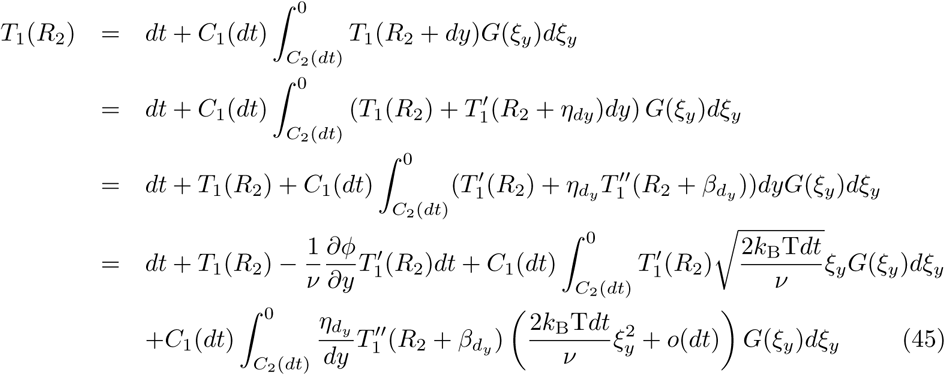

where we used mean value theorem twice to reach to Eq (45) with *R*_2_ + *dy < R*_2_ + *η*_*dy*_ *< R*_2_ + *β*_*dy*_ *< R*_2_. Note that *β*_*dy*_ depends on *η*_*dy*_ and thus depends on *dy. C*_2_(*dt*) is the value to satisfy *R*_2_ + *dy* = *R*_1_ for given *dt* and *ξ*_*y*_. *C*_1_(*dt*) is the scaling factor such that the integral of probability density equals 1: 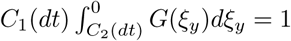where 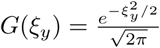. After some re-arrangements and dividing by *dt* on both sides,

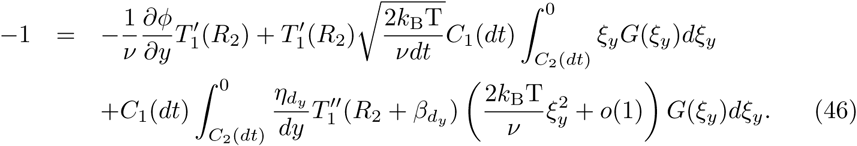

As *dt →* 0, *C*_1_ *→* 2, *C*_2_ *→ −∞*. Note that 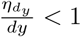 and we have 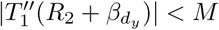 for some *M* because *T*_1_ is *C*^2^. Then if we take *t→ ∞* the third term in RHS of Eq (46) can be bounded as,

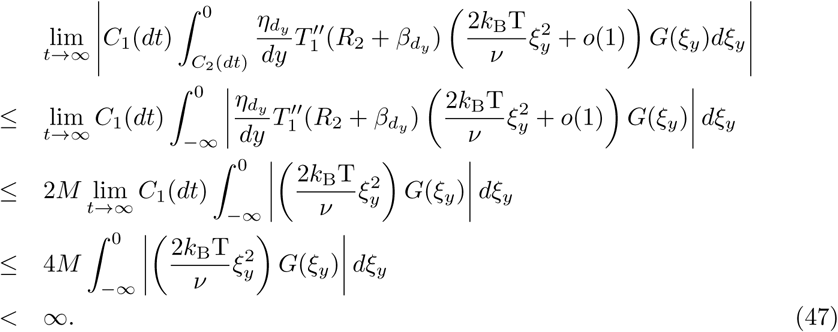

The first term in the RHS of Eq (46) is independent of *dt* and thus is finite as *t → ∞.* For the second term in RHS of Eq (46), 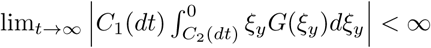, but 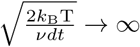 as *dt →* 0. Since the LHS of Eq (46) is finite, we must have 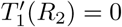.

**Proof of Theorem 2** We transform Eq (27) into polar coordinates,

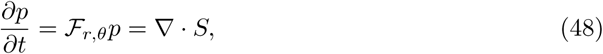

where the elliptic differential operator *ℱ*_*r,θ*_ : 𝒟(*ℱ*_*r,θ*_) *⊂ C*^2^([*R*_1_, *R*_2_] *×* [0, 2*π*]) *→ C*^2^([*R*_1_, *R*_2_] *×* [0, 2*π*]) is in divergence form, with domain

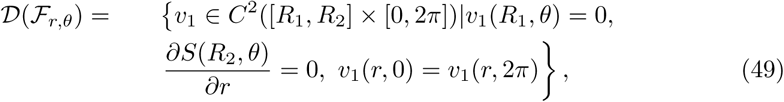

and the inner product is defined as,

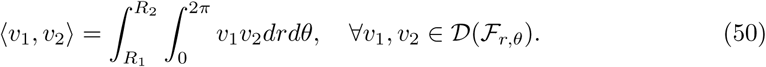

The expression of *ℱ*_*r,θ*_ can be found in [26] and we ignore the expression of *S* for brevity. Then, it’s useful to derive 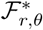 (see [26]), the adjoint operator of *ℱ*_*r,θ*_ that satisfies 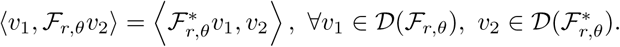.

Next, we integrate Eq (48) for *p* over *t ≥* 0 and get,

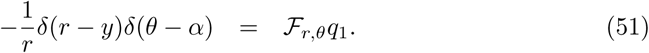

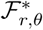 is uniformly elliptic with certain boundary conditions the solution of which has been discussed in [35]. Then, we can find a *u*_0_ *∈ C*^2^([*R*_1_, *R*_2_] *×* [0, 2*π*]) s.t. 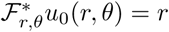. It follows from Eq (29) that,

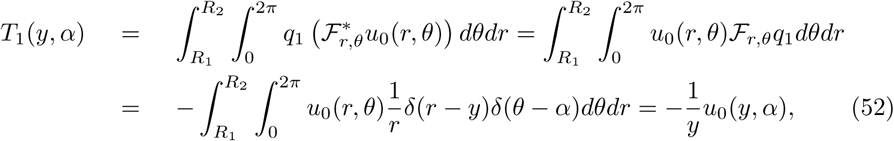

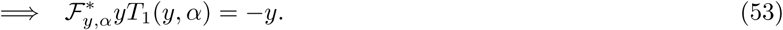

Then, we can derive a second order PDE for *T*_1_(*y*) (Eq (31)). For boundary conditions, we just need to worry about the reflecting wall. For anisotropic case, *dy <* 0. *dθ* could be either positive or negative. Similarly we can write,

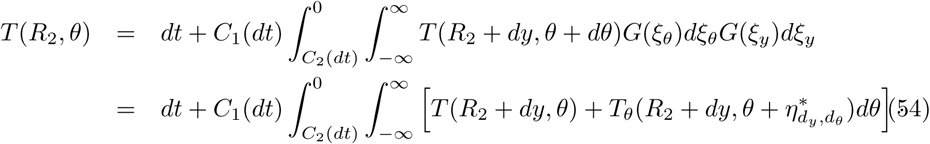

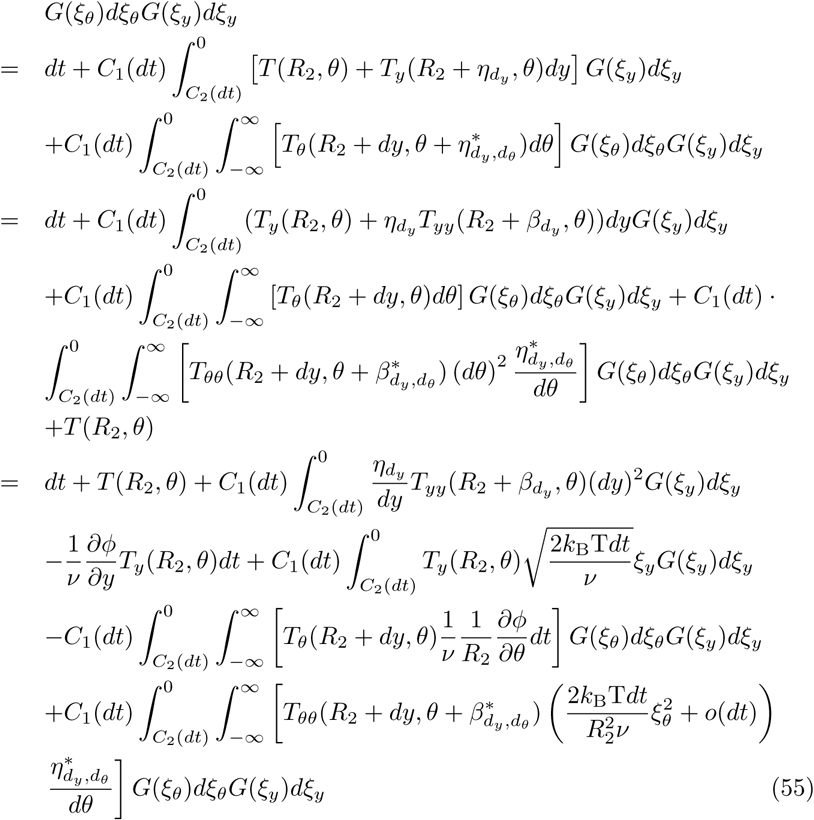

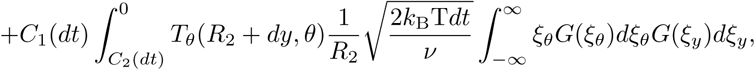

where in the process to Eq (55) we used mean value theorem three times with 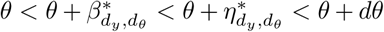 if *dθ >* 0 and 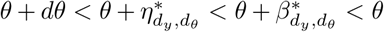 if *dθ <* 0. After some re-arrangements and dividing by *dt* on both sides, we get

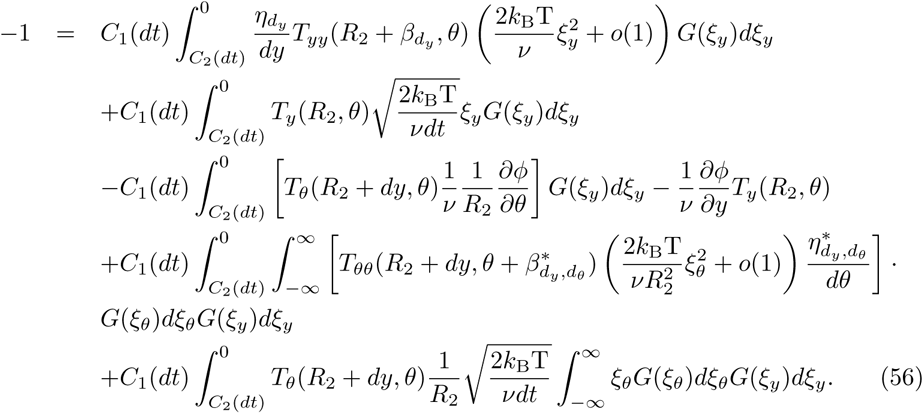

Using the continuity of *T*_*θ*_, *T*_*yy*_ and *T*_*θθ*_ and the fact that 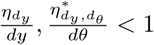, it is clear that the terms in the first, third and fourth line of Eq (56) are finite as *dt →* 0. The term in the last line is vanishing due to 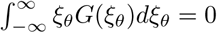. Since the LHS of Eq (56) is finite also, the term in the second line of Eq (56) must also be finite as *dt →* 0. Accordingly, *T*_*y*_(*R*_2_, *θ*) = 0 follows from 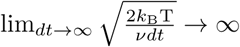.

## Acknowledgments

We acknowledge support for this work through an NSF grant CMMI 1662101.

## Footnotes

1 We are limited in the shapes we can explore by the equilateral triangle grid used in our computations.

